# The association of the RSC remodeler complex with chromatin depends on prefoldin-like Bud27 and determines nucleosome positioning and polyadenylation sites usage in *Saccharomyces cerevisiae*

**DOI:** 10.1101/2023.03.21.533665

**Authors:** A. Cuevas-Bermúdez, V. Martínez-Fernández, A.I. Garrido-Godino, A. Jordán-Pla, X. Peñate, M. Martín-Expósito, G. Gutiérrez, C.K. Govind, S. Chávez, V. Pelechano, F. Navarro

**Author notes:** A. Cuevas-Bermúdez and V. Martínez-Fernández have contributed equally to this work. Laboratoire de Biologie Moléculaire et Cellulaire des Eucaryotes, UMR8226, Centre National de la Recherche Scientifique (CNRS), Institut de Biologie Physico-Chimique, Paris Sciences et Lettres (PSL) Research University, Sorbonne Université, F-75005 Paris, France.

## Abstract

The tripartite interaction between the chromatin remodeler complex RSC, RNA polymerase subunit Rpb5 and prefoldin-like Bud27 is necessary for proper RNA pol II elongation. Indeed lack of Bud27 alters this association and affects transcription elongation. This work investigates the consequences of lack of Bud27 on the chromatin association of RSC and RNA pol II, and on nucleosome positioning. Our results demonstrate that RSC binds chromatin to gene bodies and lack of Bud27 alters this association, mainly around polyA sites. This alteration impacts chromatin organization and leads to the accumulation of RNA pol II molecules around polyA sites, likely to be arrested or slower. Our data suggest that RSC is necessary to maintain chromatin organization around those sites, and any alteration of this organization results in the widespread use of alternative polyA sites. Finally, we also find a similar molecular phenotype that occurs upon TOR inhibition with rapamycin, which suggests that alternative polyadenylation observed upon TOR inhibition is likely Bud27-dependent.

## INTRODUCTION

RNA polymerase II is the enzyme that transcribes all mRNAs and many non-coding RNAs (1). This complex comprises 12 subunits, of which the Rpb4/7 dimer is dissociable in *Saccharomyces cerevisiae* (2,3). RNA pol II transcription is divided into three major phases, that correspond to the formation of the PIC (pre-initiation complex), elongation (after promotor scape of RNA pol II) and termination (comprising polyadenylation, primary transcript cleavage and enzyme release) (4).

Transcription elongation by RNA pol II is linked to some post-transcriptional processes such as splicing or polyadenylation (5–8). Elongating RNA pol II competes with the machinery involved in cleavage/polyadenylation, and affects poly(A) sites (pAS) selection (7). It has been reported that mutants affected in transcription elongation, such as those showing defects in Rpb2, and transcriptional elongation factors TFIIS and Spt5, affect alternative polyadenylation in *S. cerevisiae* (9). Furthermore, the RNA pol II elongation rate is associated with alternative polyadenylation in response to environmental conditions in yeast (7), a link that has also been established in mammals and *Drosophila* (10,11).

During transcription elongation, different RNA pol II subunits, such as Rpb5 and Rpb4/7, as well as distinct elongation factors, such as TFIIS an Spt4/5, among others, are necessary (2,12–16). Furthermore, the post-translational modifications of the CTD domain of Rpb1, the largest RNA pol II subunit, are essential for this process, being the most relevant the CTD-Ser2 phosphorylation (17,18).

RNA pol II elongation is influenced by chromatin structure, and then by nucleosomal organisation and histone modifications, and *vice versa* (19,20). The association between chromatin alteration and alternative polyadenylation has been previously reported (21–23) and is likely related to epigenetic modifications, such as H3K36me3, H3K4me1 and H3K4me3 (21–24).

Cells use chromatin remodeler enzymes to act on chromatin and to regulate gene expression. Chromatin remodelers invest the energy of ATP to assemble, evict, slide or restructure nucleosomes (25,26). One of these remodelers is the RSC (Remodels the Structure of Chromatin) complex, a member of the SWI/SNF family and the only essential member in budding yeast (27). RSC has been implicated mainly in transcription initiation by establishing ‘nucleosome-depleted regions’ (NDRs) upstream of transcription start sites (TSSs) and by positioning NDRs-flanking nucleosomes (the nucleosomes −1 and +1) and, to a lesser extent, the nucleosomes +2 and +3 (28). In addition to its role in transcription initiation, RSC has been reported to act in transcription elongation both *in vitro* (29) and *in vivo* (30–32) and has been suggested to also influence transcription termination (32).

RSC contacts RNA pols through its Rsc4 subunit and the Rpb5 subunit that is common to RNA pols I, II and III (33). The existence of a tripartite interaction between RSC-Rpb5 and prefoldin-like Bud27 in *S. cerevisiae* has been proposed, which is necessary for RNA pol II transcription elongation (34). Lack of Bud27 affects the association of RSC with chromatin and RNA pol II, and alters elongation (34 Notably, a functional connection between RSC and Bud27, which influences RNA pol III transcription, has been reported {Vernekar, 2015 #1421).

Bud27 and its human orthologue URI (unconventional prefoldin RPB5-interactor) are members of the prefoldin family of ATP-independent molecular chaperones, which function as scaffold proteins during the assembly of additional prefoldin family members (35,36). Both Bud27 and URI bind the Rpb5 subunit of the three RNA pols (35,37,38). Bud27 stimulates not only the transcription elongation by RNA pol II (34), but also the transcription of ribosomal components by the three RNA pols (39), and URI also participates in transcription (38,40–43). Bud27 and URI play roles in different cellular processes; for example in: the assembly of RNA pols (44–47); TOR-dependent transcription programmes (35,37,40,45,48) and translation initiation and cotranslational quality control (49). In addition, human URI is related to cancer development (50–52).

In this work, we use a mutant of *S. cerevisiae* that lacks prefoldin-like Bud27, which alters the association of the chromatin RSC remodeler complex with both RNA pol II and chromatin, and affects transcriptional elongation. We demonstrate that RSC associates with chromatin along the gene bodies and lack of Bud27 affects this association through the 3’ region, influences nucleosome positioning and the chromatin structure around pAS, and increases the likely paused or slower RNA pol II. Consequently, this alteration affects pAS usage and, thus, alternative polyadenylation. We also demonstrate that TOR inhibition by rapamycin addition leads to similar behaviour than the absence of Bud27, indicating that the impact of TOR inhibition on polyadenylation is likely Bud27-dependent.

## MATERIALS AND METHODS

### Yeast strains and growth conditions

The *Saccharomyces cerevisiae* strains used in this work are BY4741 and YFN105, along with their derivatives YFN360 and YFN359 containing the *STH1::13MYC::Kan-MX6* allele (45).

Media preparation, yeast transformation and genetic manipulations were used as described elsewhere (53). Yeast strains were cultured in YPD at 30 °C.

The rapamycin-treated cells corresponded to the cells grown in YPD at 30°C, which contained 50 ng/mL of rapamycin until an OD600 of about 0.8

In liquid cultures, cell growth was constantly monitored to maintain them in exponential phase.

### Chromatin immunoprecipitation

The chromatin immunoprecipitation experiments were performed using magnetic beads Dynabeads M-280 sheep anti-mouse IgG (Invitrogen), mixed with an anti 9E10 C-myc antibody (Santa Cruz), or Dynabeads M-280 Sheep Anti-rabbit IgG mixed with anti-anti-Rpb1 (39) or anti-Histone H3 (ab1791; Abcam) antibodies, as previously described (54). For real-time PCR, 1:100 dilution was used for the input DNA and 1:4 dilution for the DNA of the immunoprecipitated samples. Genes were analysed by quantitative real-time PCR in triplicate with at least three independent biological replicates using SYBR premix EX Taq (Takara). Quantification was performed as indicated in the figure legends. The oligonucleotides utilised for the different PCRs are listed in Supplemental Table S1.

### Chromatin-enriched fractions, SDS-PAGE, western blot analysis and immunoreactive bands quantification

The chromatin-enriched fractions were prepared by following the yChEFs procedure (55) from 50 ml of the yeast cells grown in YPD at 30 °C until the mid-log phase.

Protein electrophoresis and western blot were carried out as described in (54). The western blots of the chromatin-associated proteins were performed with anti-RNA pol II CTD phospho Tyr1 (Active Motif), anti-Rpb1 (manufactured in our laboratory) (39), anti-phosphoglycerate kinase, Pgk1, (459250, Invitrogen) and anti-H3 (ab1791; Abcam). The intensities of the immunoreactive bands on the western blots were quantified by densitometry using the IMAGE STUDIO LITE software from the images acquired with an office scanner.

### RNA extraction and reverse transcription

Total RNA from yeast cells was prepared and quantified as previously described from 50 ml cultures, which were grown in appropriate medium until an OD600 of about 0.8.(54). First-strand cDNA was prepared from 0.5 μg of RNA with the iScript cDNA synthesis kit (Bio-Rad) following the manufacturer’s protocol. Each sample was subjected to the same reaction without reverse transcriptase, as a negative control for genomic DNA contamination.

For the RNA analysis by real-time PCR, a CFX-384 Real-Time PCR instrument (BioRad) and the the EvaGreen detection system ‘‘SsoFast™ EvaGreen® Supermix’’ (BioRad) were used. Reactions were performed in a 5 µl volume with 1:100 dilution of the synthesised cDNA. Each sample was analysed in triplicate with at least three independent biological replicates per sample. Values were normalised to the steady-state 18S rRNA levels. The used oligonucleotides are listed in Supplemental Table S1.

### mRNA stability analysis

For the mRNA stability analysis, cells were grown in SD (with requirements) to an OD_600_ ∼ 0.5. At that time, cells were treated with 5 μg/ml thiolutin. Next 15-mL aliquots of cell samples were taken at different times after thiolutin addition (up to 120 min), and were then pelleted and frozen. Total RNA was isolated from these samples and cDNA was synthesised as previously described (54). The mRNA stability (half-lives) for the selected genes was analysed following the decay curves obtained by RT-qPCR with specific primers for the corresponding genes (see Table S1). Similarly was proceeded for global mRNA decay analysis, with 5 μg of RNA, by dot-blot using a fluorescent oligo-dT probe, as previously described (56).

### Nucleosome mapping with MNase

The MNase digestion was performed as described in (57). Briefly, 500 ml of exponentially grown yeast culture in YPD medium was cross-linked with formaldehyde. Cells were washed and resuspended in Buffer Z2 (1 M Sorbitol; 50 mM Tris-HCl pH 7.4; 10 mM β-mercaptoethanol). Zymolyase was added and cells were incubated for 30 min at 30 °C to obtain spheroplasts. Spheroplasts were incubated with different amounts of micrococcal nuclease (from 500 to 10 mU). Protein was degraded by Proteinase K and DNA was obtained by phenol-chloroform-isoamyl extraction and ethanol precipitation. Digested DNA was resolved on an agarose gel and the band corresponding to mononucleosomes was purified. Naked DNA was obtained as before, but MNase was added after DNA extraction instead of before, and a band with similar digestion as the mononucleosomes was purified.

The MNase-digested mononucleosomal fragments of DNA that resulted from the wild-type and *bud27Δ* strains and the corresponding nucleosome-sized fragments of naked DNA were sent to the genomic facility of the Centre of Genomic Regulation (CRG) in Barcelona to be sequenced. These DNA fragments were sequenced by the paired-end technology from Illumina (HiSeq Sequencing v4 Chemistry with read length 50).

After QC v0.9.6 (58) was used with default parameters to do the quality control, the filtering and trimming of the paired-end sequencing data obtained from MNase were performed. The cleaned sequence data were aligned to the *S. cerevisiae* (R64) reference genome with bowtie2-2.3.3.1 (59). Once mapped, the resulting SAM files were analysed using DANPOS v2.2.2 (60) to determine nucleosome localisation, occupancy and fuzziness across the whole genome in all the analysed strains. DANPOS pools the replica of the same condition and can use naked data as background correction.

The accession number for the data corresponding to the *bud27Δ* strains is GSE226720. As a wt reference, we used previously published data with accession number GSE153037.

### Bioinformatic analyses

For the Rpb1 ChIP-Seq analysis, previously reported datasets were used (GEO, accession number GSE131390, (39)).

The polyadenylation analysis was carried out by using a previously published dataset for RNA-Seq (GEO, accession number GSE131390, (39)), which was prepared with the QuantSeq 3’ mRNA-Seq Library Prep Kit from Illumina following the manufacturer’s instructions. The 3’T-fill method that allows sequencing to start immediately after de polyA tail was applied (57).

For the Sth1-Myc ChIP-Seq analysis, paired-end ChIP-sequencing datasets for the two biological replicates of Sth1-Myc IP and the two biological replicates of their corresponding input DNAs were downloaded from EBI ENA (PRJNA612872, (31)) and mapped to the *S. cerevisiae* reference genome (*SacCer3*) with bowtie2 (v2.4.4) (59) and the following settings: -X 1000 --very-sensitive --no-mixed --no-unal. No pre-processing steps were necessary given the high quality of the reads in the public datasets. Next, replicate similarity was assessed and, due to the high degree of concordance, the two replicates of each sample type were merged into a single alignment file with the merge function of Samtools (v1.12) (61). Merging resulted in one IP and one input dataset, which we then used for the downstream analyses.

For the metagene plots, the normalised average density plots around genomic features were calculated with the R and Python software (ngs.plot, v2.61) (62) using indexed alignment files as inputs and the internal *SacCer3* database annotation as a reference. The statistical robustness parameter, which filters out 0.5% of genes with the most extreme (high and low) count values, was applied to all the calculations. The fragment length parameter was set at -FL 300 and at -FL 10 for the ChIP-Seq and pA mapping datasets, respectively.

## RESULTS

### Lack of Bud27 affects Sth1 occupancy through the 3’ region of genes

Absence of Bud27 affects RNA pol II transcription elongation by altering the association of the RSC remodeling complex with RNA pol II (39).

We wondered whether this defect could affect RSC occupancy along transcription units. To do so, we analysed Sth1 occupancy (the catalytic subunit of the RSC remodeling complex), by chromatin immunoprecipitation (ChIP-qPCR), in the wild-type and *bud27Δ* mutant strains that contain a functional Myc-tagged version of Sth1 (39). According to previously published data of mRNA expression (39), we selected some up-regulated (*CPA2* and *IRC24*) and down-regulated (*TEF2* and *PYK1*) genes to analyse occupancy along their transcription units in *bud27Δ vs.* the wild-type strain (mRNA levels were corroborated by RT-qPCR; data not shown) at 30°C. As shown in Figure 1A, Sth1 occupancy decreased throughout the 3’ region of all the analysed genes in the *bud27Δ* strain, albeit to a different extent, and no differences were observed for the 5’ regions.

**Figure 1:**
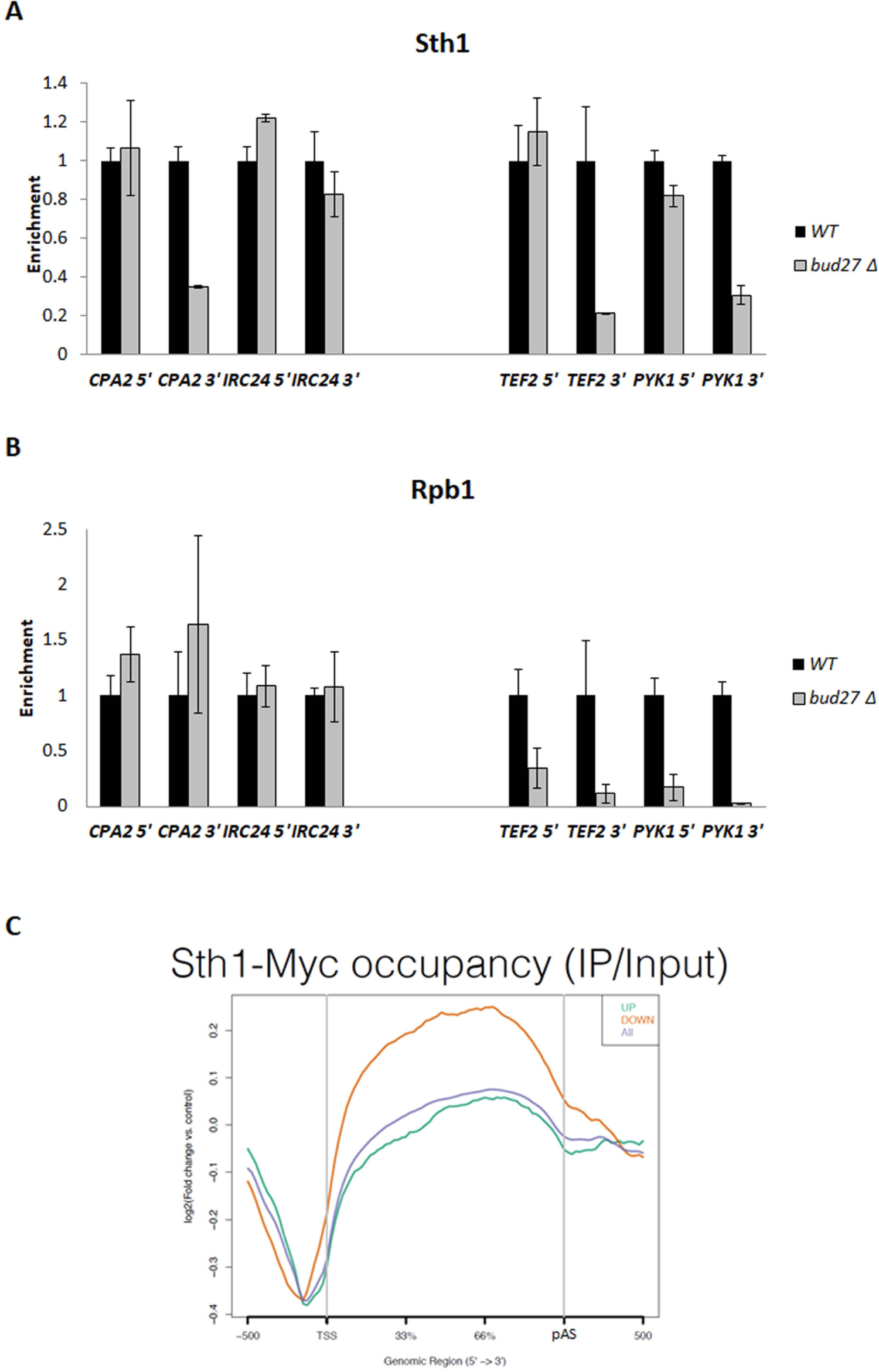
Lack of Bud27 affects Sth1 occupancy. Chromatin immunoprecipitation (ChIP) analysis for different genes in wild-type (WT) and *bud27Δ* cells, performed with anti-Myc (**A**) and anti-Rpb1 (**B**) antibodies, against Sth1-Myc and the Rpb1 subunit of RNA pol II. The values found for the immunoprecipitated PCR products were compared to those of total input, and the ratio of each PCR product of transcribed genes to a non-transcribed region of chromosome V was calculated. The values obtained in the wild-type cells were represented as 1 for all the tested genes. **C**) Metagene analysis of Sth1 occupancy, in a wild-type strain, by using the RSC MNase ChIP-Seq data from (31).

We also analysed Rpb1 occupancy (Figure 1B). Unlike RSC, and in agreement with mRNA levels, Rpb1 occupancy drastically decreased in both the 5’ and 3’ regions for the down-regulated genes, while it did not seem to be altered for the tested up-regulated genes.

We re-analysed the previously reported genome-wide dataset of RSC MNase ChIP-Seq in a wild-type strain (31) to evaluate Sth1 occupancy in relation to the up and down-regulated genes observed in the *bud27Δ* mutant (39). Interestingly, as Figure 1C depicts, the down-regulated genes in the *bud27Δ* mutant corresponded to those showing greater Sth1 occupancy in a wild-type strain. These results suggest that lack of Bud27 preferentially affects the expression of the most Sth1-occupied genes.

### Lack of Bud27 alters nucleosome positioning in gene bodies and affects chromatin structure around polyA sites

It has been reported that the RSC remodeling complex plays a role in transcription initiation and elongation (19,28,29,31,32,63,64). In addition, RSC depletion alters nucleosome positioning *in vivo* (28,31,32,63–65).

Based on that evidence, we wondered whether lack of Bud27 could also alter nucleosome positioning. To do so, we performed MNase-Seq by following a refined nucleosome mapping method in *S. cerevisiae* (66) and analysed the global nucleosomal changes caused by lack of Bud27 in the *bud27Δ* mutant in relation to its isogenic wild-type strain, at the permissive temperature of 30°C. As Figure 2A illustrates, a general increase in nucleosome positioning along transcription units was observed in *bud27Δ* in relation to a wild-type strain for all genes, which is likely associated with a sharp drop in RNA pol II-dependent transcription in this mutant (39). These results suggest a correlation between nucleosome occupancy and transcriptional activity in *bud27Δ*, as shown for the increase in nucleosome positioning for the down-regulated genes *versus* the up-regulated ones (Supplemental Figure S1A). This behaviour was corroborated by the high nucleosome positioning for RP genes which showed lower mRNA expression in *bud27Δ* cells (39) (Supplemental Figure S2A).

**Figure 2:**
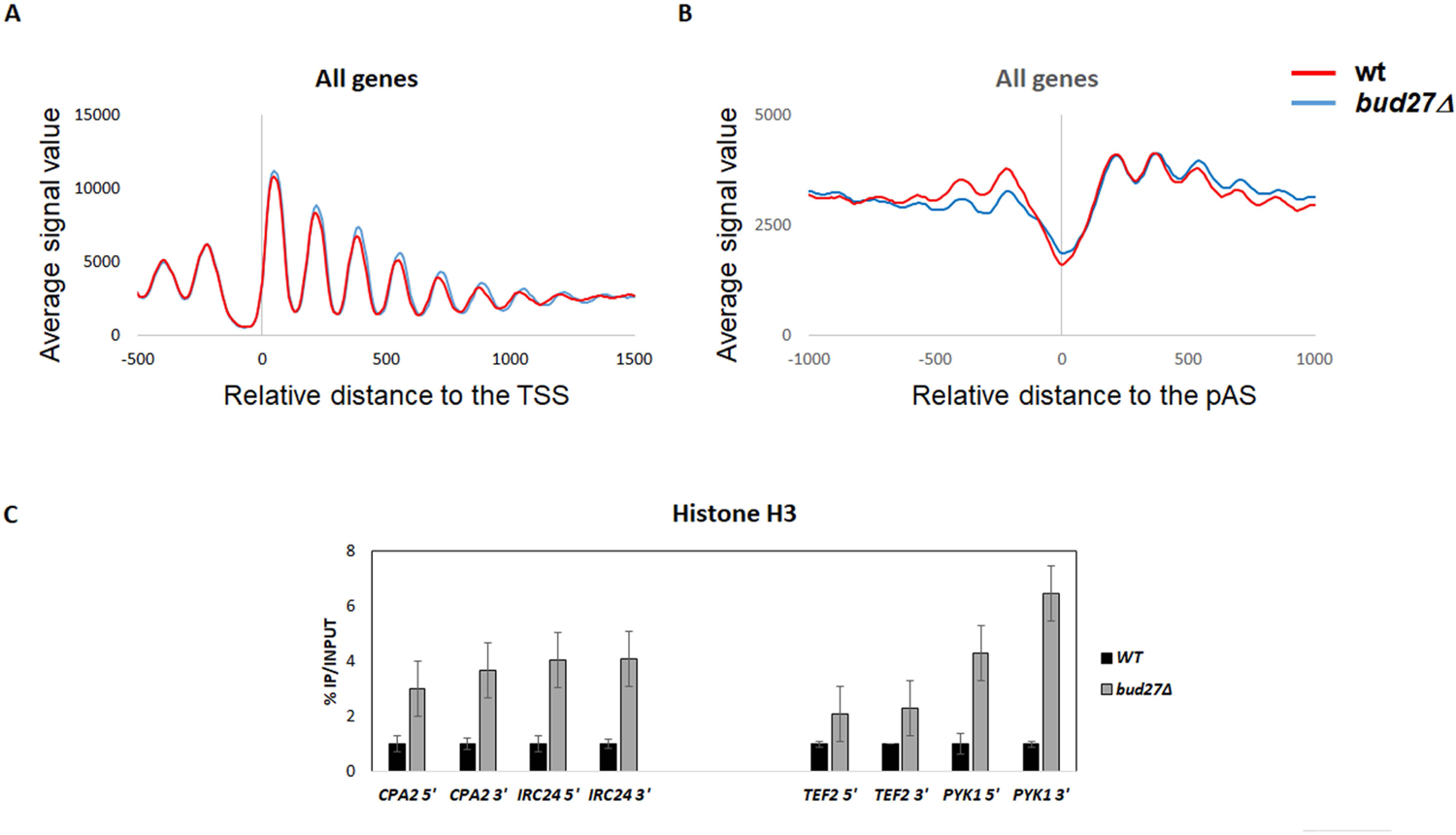
Lack of Bud27 alters nucleosome positioning. **A**) and **B**) Metagene analysis of nucleosome occupancy for wild-type (red) and *bud27Δ* cells (blue). Genes were scaled to the same length and then aligned to their TSS (**A**) or their pAS (**B**). All the genes in the yeast genome, for which a TSS/pAS is available, were considered. **C**) Chromatin immunoprecipitation (ChIP) analysis for different genes in the wild-type and *bud27Δ* cells performed with an anti-H3 antibody, against histone H3. The values found for immunoprecipitated PCR products were compared as % of immunoprecipitated material, IP, *vs,* INPUT. The values in the wild-type cells were represented as 1 for all the tested genes.

Furthermore, the lack of Bud27 led to an overall alteration in nucleosome positioning along gene bodies, which consisted of a clear shift towards 3’ (Figure 2). Such behaviour was observed for both the up and down-regulated genes in *bud27Δ* cells (Supplemental Figure S1).

According to the information above, and as lack of Bud27 decreased Sth1 occupancy through the 3’ region of genes, we next checked whether chromatin structure could be altered around pAS in *bud27Δ* mutant cells. To do so, we analysed the nucleosome positioning around pAS in the *bud27Δ* mutant and its isogenic wild-type strain. As seen in Figure 2B, a clear alteration of chromatin structure around pAS was observed in *bud27Δ* cells with respect to wild-type cells (Note that observation in yeast showed less defined nucleosomal profiles around pAS than around the transcription start site (TSS), which correlates with the presence of AT rich regions in the vicinity of pAS (66)). This effect was particularly evident for the most highly expressed genes and less marked for the down-regulated genes in *bud27Δ* cells (Supplementary Figures S1B and S2B). This scenario suggests a correlation between chromatin structure around pAS and transcriptional activity in *bud27Δ*.

Histones are the constituents of nucleosomes, and histone density increases at RNA pol III loci with Sth1 protein loss (67). Then we analysed histone H3 occupancy by ChIP at the 5’ and 3’ regions of gene bodies for several RNA pol II genes. As shown in Figure 2C, histone H3 occupancy increased at all the analysed RNA pol II genes. These results agree with what has been previously reported at the RNA pol III locus upon Sth1 loss (67).

Taken altogether, these data demonstrate that lack of Bud27 affects chromatin structure at the RNA pol II locus by altering nucleosome positioning and occupancy along gene bodies and around pAS, likely as a result of both deficiency in the RSC remodelling complex association with chromatin and the defect in transcription elongation.

### *bud27Δ* cells alter polyadenylation patterns and shift to uptream polyA site selection

The transcription elongation rate is associated with alternative polyadenylation in yeast and other organisms (7,9–11). In addition, some evidence has been reported about the relation between chromatin structure and alternative polyadenylation (21–23).

We wondered whether the altered chromatin organisation around pAS provoked by a defect in RSC-RNA pol II association in *bud27Δ* cells (34) and the decreased RSC binding throughout the 3’ region of genes could affect polyadenylation site selection. To this end, we further characterised genome-wide the effect of lack of Bud27 on pAS selection at 30°C by following the 3’T-fill method, which allows sequencing to start immediately after the polyA tail and to quantify 3’ polyadenylation isoforms (57). By this methodology, each polyA site call is defined as the read base closest to the polyA tail (the first base in the 3’T-fill protocol).

As shown in Figure 3A, the profile of pAS selection over 150 nt upstream and downstream of pAS, suggested that it was wider in *bud27Δ* cells *versus* the wild-type strain, at least for the region corresponding to −50 / +50 nt around pAS, and bigger upstream the canonical pAS. Interestingly, and as previously shown for nucleosome accumulation, the widest variation in pAS selection correlated with transcriptional activity, and was maximal for the up-regulated genes in *bud27Δ* cells (Supplemental Figure 3). Note that the same was true for the wild-type strain because the most highly expressed genes corresponded to the down-regulated genes in *bud27Δ* cells (Supplemental Figure 3).

**Figure 3:**
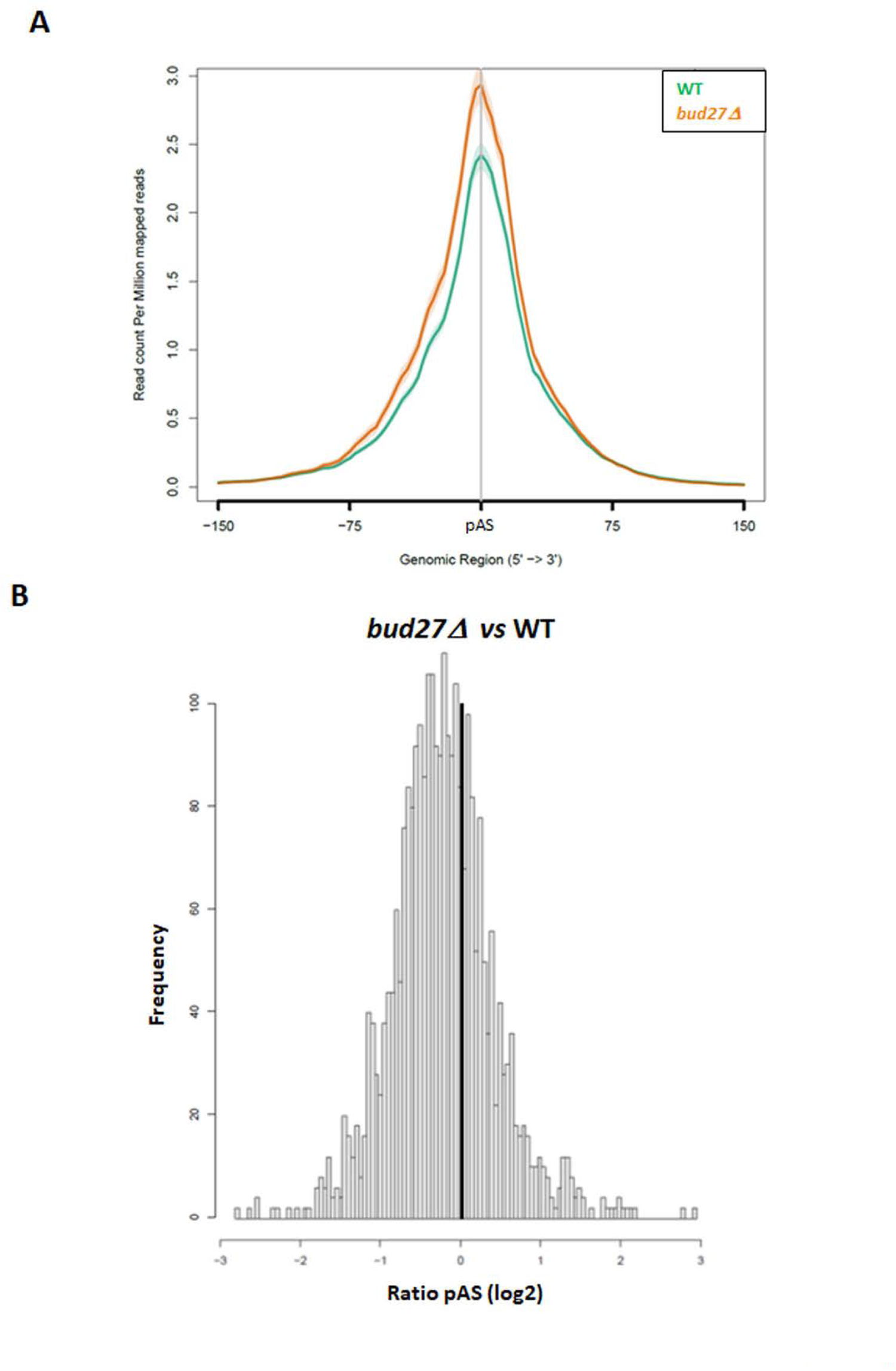
Polyadenylation pattern is altered in *bud27Δ* cells and shift to upstream polyA site selection. **A**) Metagen analysis from RNA-Seq data represented as read count per million mapped reads in wild-type (WT) and *bud27Δ* cells in the −150 to +150 region around the canonical annotated pA site. **B**) Gene-specific frequency to use preferentially pAS upstream or downstream of the canonical annotated pAS. For each gene, we firstly computed its preferential use of downstream or upstream pAS (e.g., Log_2_(reads_down_/ reads_up_)). Next, we compared the different strains. The X-axis shows the ratio Log_2_(reads *bud27Δ*_down_ / reads *bud27Δ*_up_) - Log_2_(reads WT _down_/ reads *bud27Δ*_up_), where a negative value represents preferential upstream pAS use in the mutant. The Y-axis shows the frequency of the genes corresponding to each value in X for *bud27Δ vs.* the wild-type (WT) cells.

We analysed if there was a shift in pAS usage in *bud27Δ* cells. To do so, for all the genes we determined the polyadenylation sites usage at a single nucleotide resolution, and both upstream and downstream of the canonical pAS. Then we represented them as the ratio by comparing pAS usage in the *bud27Δ* mutant to its isogenic wild-type strain. As Figure 3B depicts, the *bud27Δ* mutant showed a preference for upstream pAS usage in relation to those used in the wild-type strain.

Our data collectively demonstrate that the mutant lacking Bud27 affects alternative polyadenylation and favoured the use of the upstream pAS. This behaviour likely correlates with the chromatin structure alteration around pAS that resulted from deficiency in the RSC remodelling complex. In agreement with this, the RNA pol II mutants with low elongation rates shifted to the upstream pAS selection, while a shift to the downstream pAS occurred in the mutants with high elongation rates (7).

### Transcription elongation defects and chromatin structure alteration are accompanied by alterations in RNA pol II occupancy in *bud27Δ* cells

Based on the results above and previously reported data (39), we wondered whether transcription elongation defects and chromatin structure alteration in *bud27Δ* cells could correlate with RNA pol II occupancy. For this purpose we profiled global Rpb1 occupancy along transcription units, by using previously published data from ChIP-Seq in wild-type and *bud27Δ* mutant strains (39).

As shown in Figure 4A, an overall increase in Rpb1 occupancy took place in the *bud27Δ* mutant in relation to the wild-type strain in the gene bodies for all the groups of genes. Note that this increase was maximal for the up-regulated genes and correlated with greater transcriptional activity. Notably from at least 2/3 of the gene body to the 3’ region, the Rpb1 accumulation pattern differed for the *bud27Δ* mutant in relation to the wild-type strain. In the wild-type strain, Rpb1 significantly increased in the last 1/3 of the gene body to pAS, and this increase was moderate for the *bud27Δ* mutant strain. Moreover, the data from Figure 4A (left panel) suggest diminished Rpb1 accumulation around pAS in the *bud27Δ* mutant compared to the wild-type strain when all the genes were analysed together. To gain better insight, we focused on the regions around pAS and analysed Rpb1 occupancy in the −500 / + 500 nt regions from pAS (Figure 4B), which corroborated the above-indicated results. These data suggest slower or arrested RNA pol II, likely by the deficiency in the RSC association and chromatin structure alteration. In addition, the decrease in Rpb1 accumulation downstream of pAS that was observed in the wild-type strain (Figure 4B) was altered in the *bud27Δ* mutant strain which showed a less precipitous reduction in Rpb1 occupancy. According to these results, we cannot rule out a defect in transcription termination, although the analysis of the Tyr1 phosphorylation of RNA pol II, a mark of transcription termination, did not reveal any significant differences between the *bud27Δ* mutant and the wild-type strain (Figure S4).

**Figure 4:**
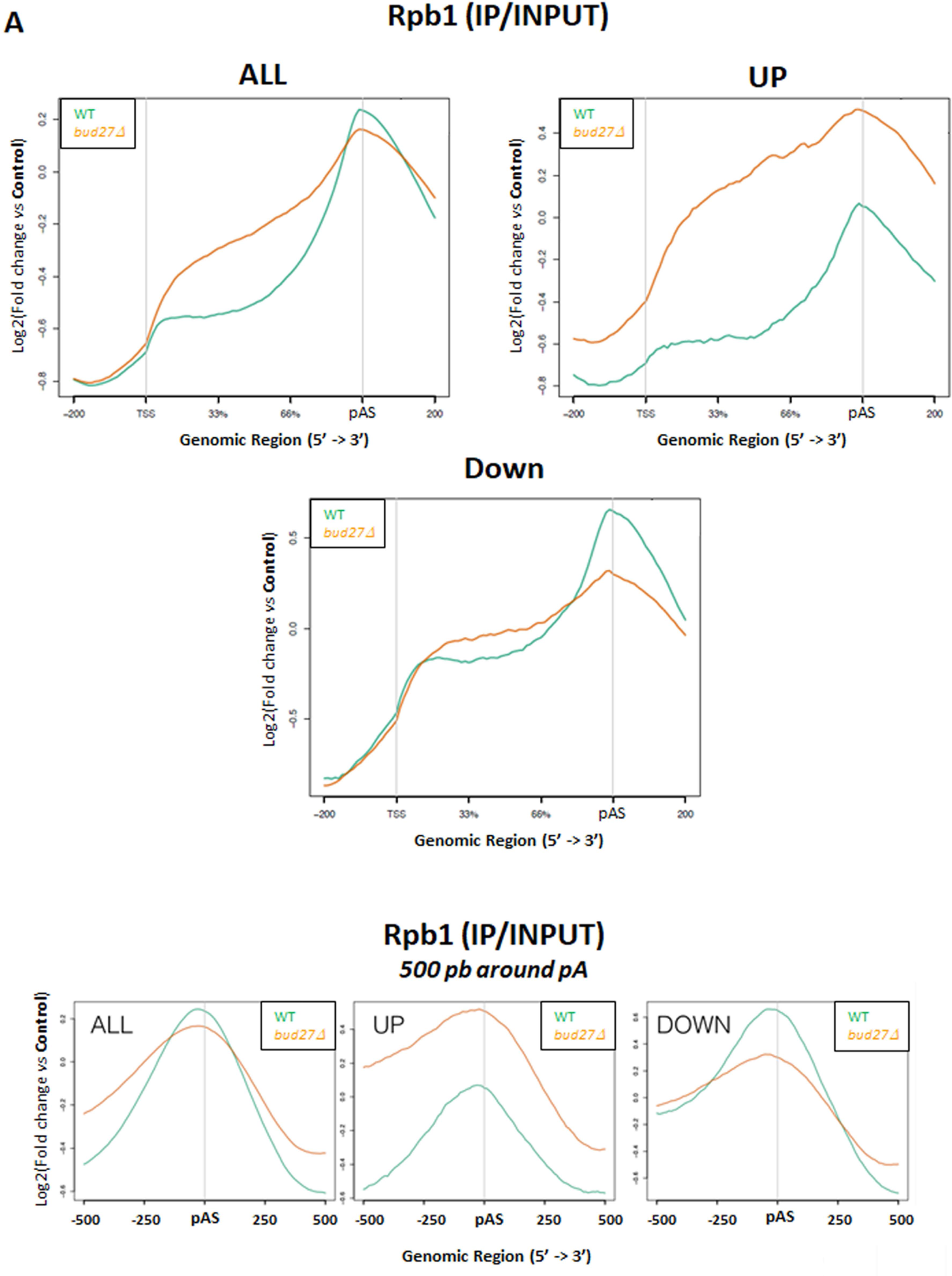
Rpb1 occupancy. **A**) Metagene analysis of Rpb1 occupancy along the transcription units in the wild-type and *bud27Δ* mutant strains for all the genes (ALL), or those which are up or down-regulated in the *bud27Δ* mutant *vs.* the wild-type (WT) according to (39). **B**) As in **A** for Rpb1 occupancy around pAS.

All these data collectively indicate that lack of Bud27 affects Rpb1 occupancy along transcription units, and likely represents slower or arrested RNA pol II, which results from deficiency in the RSC association and chromatin structure alteration, which is maximal at pAS.

### Effects of blocking the TOR signaling cascade on chromatin structure and polyA site selection could, at least partially, depend on Bud27

Bud27 and its human orthologue URI (unconventional prefoldin RPB5-interactor) have been shown to participate in TOR-dependent transcription programmes (35,37,39,40), and lack of Bud27 partially mimics TOR signaling cascade inhibition (39,48). Interestingly, blocking the TOR signaling cascade by rapamycin addition alters chromatin structure around pAS and favors upstream pAS usage (22).

To explore if a correlation exists between the data shown herein for the *bud27Δ* and TOR pathway inhibition, we first analysed Sth1-myc occupancy in the wild-type yeast cells containing a functional Myc-tagged version of the Sth1 grown in YPD medium at 30°C in both the absence and presence of rapamycin for 12 h to inactivate the TOR pathway. As Figure 5A depicts, decreased Sth1-myc occupancy was observed upon rapamycin addition, with the maximal in the 3’ region of most tested genes. These results indicate a similarity between the Sth1 occupancy that resulted from TOR inhibition by rapamycin and the Sth1 occupancy alteration provoked by lack of Bud27 (the above results). In addition, we observed similar Rpb1 occupancy that was observed in *bud27Δ* mutant (Figure 5B), with no significant variations along the transcription unit. These results suggest that the decreased RSC occupancy which resulted from TOR signalling cascade inhibition could account for the alteration to chromatin structure around pAS, which has been previously reported for rapamycin treatment (22), and could influence nucleosome positioning. Therefore, we measured histone H3 occupancy by ChIP in the 5’ and 3’ regions of the gene bodies for several RNA pol II genes in cells grown in YPD medium at 30°C in both the absence and presence of rapamycin for 12 h to inactivate the TOR pathway. As seen in Figure 5C, histone H3 occupancy increased in all the analysed genes, which indicates that nucleosome occupancy must be altered and falls in line with the results observed in the *bud27Δ* mutant.

**Figure 5:**
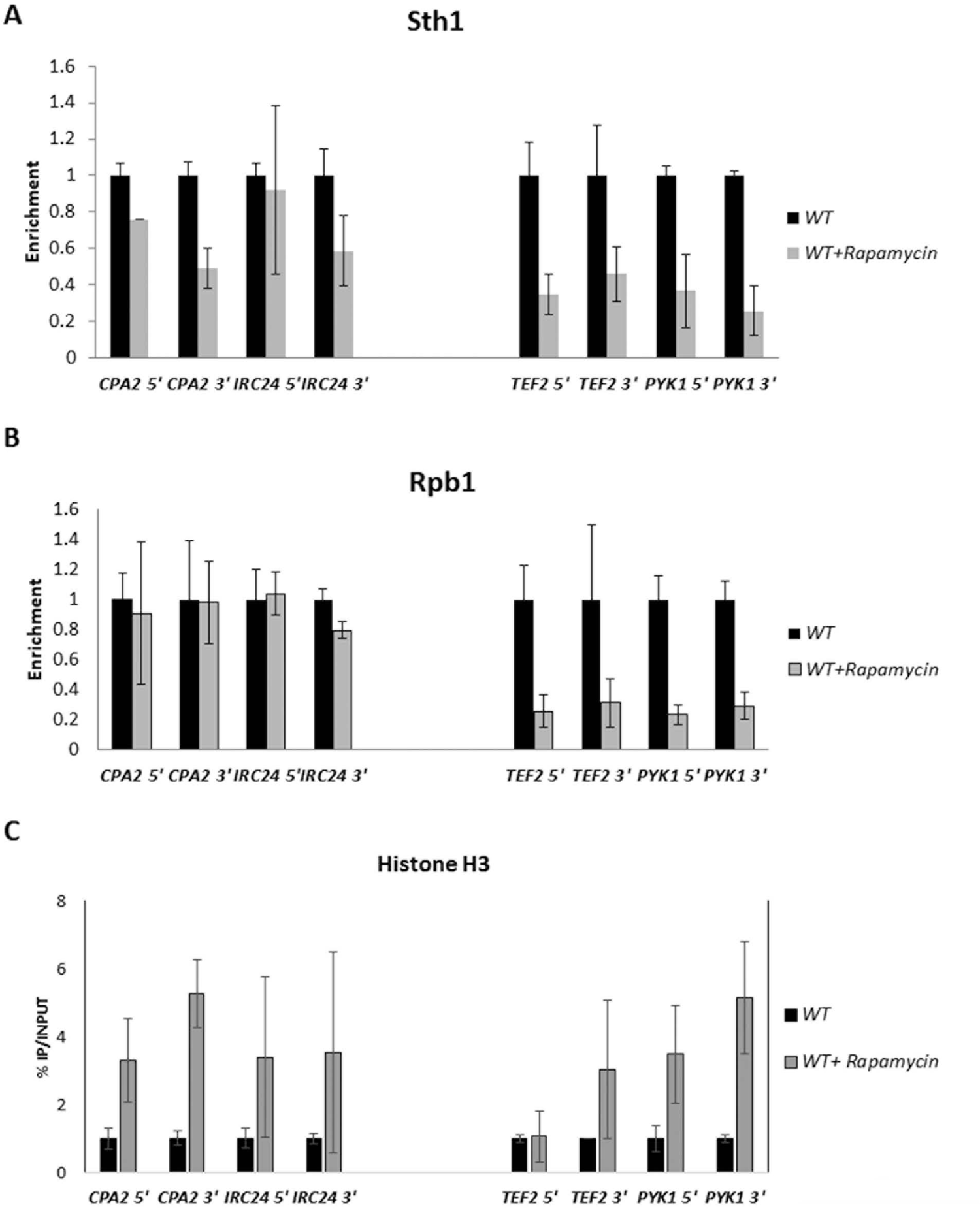
Inactivation of the TOR signaling cascade affects Sth1 occupancy. Chromatin immunoprecipitation (ChIP) analysis for different genes in the wild-type (WT) cells grown at 30°C in both the absence and presence of rapamycin for 12 h to inactivate the TOR pathway, performed with anti-Myc (**A**) anti-Rpb1 (**B**) and anti-H3 (**C**) antibodies against Sth1-Myc, the Rpb1 subunit of RNA pol II and histone H3. In **A** and **B,** the values found for immunoprecipitated PCR products were compared to those of the total input, and the ratio of each PCR product of the transcribed genes to a non-transcribed region of chromosome V was calculated. In **C**, the values found for the immunoprecipitated PCR products were compared as % of the immunoprecipitated material, IP, *vs* INPUT. The values in the wild-type cells were represented as 1 for all the tested genes.

As we demonstrated above a relationship between chromatin structure alteration in *bud27Δ* mutant and pAS selection, we wondered whether that would also be the case for rapamycin-treated cells. To investigate that we also analysed genome-wide the effect of blocking the TOR cascade on pAS selection by adding rapamycin to wild-type cells for 12 h at 30°C as indicated above. Figure 6 reveals an overall shift towards preferential upstream pAS selection use upon rapamycin treatment, which was similar to that which resulted from lack of Bud27. Furthermore, these data agree with previous results showing that yeast cells grown in the presence of rapamycin shifted to upstream pAS for some analysed genes (22).

**Figure 6:**
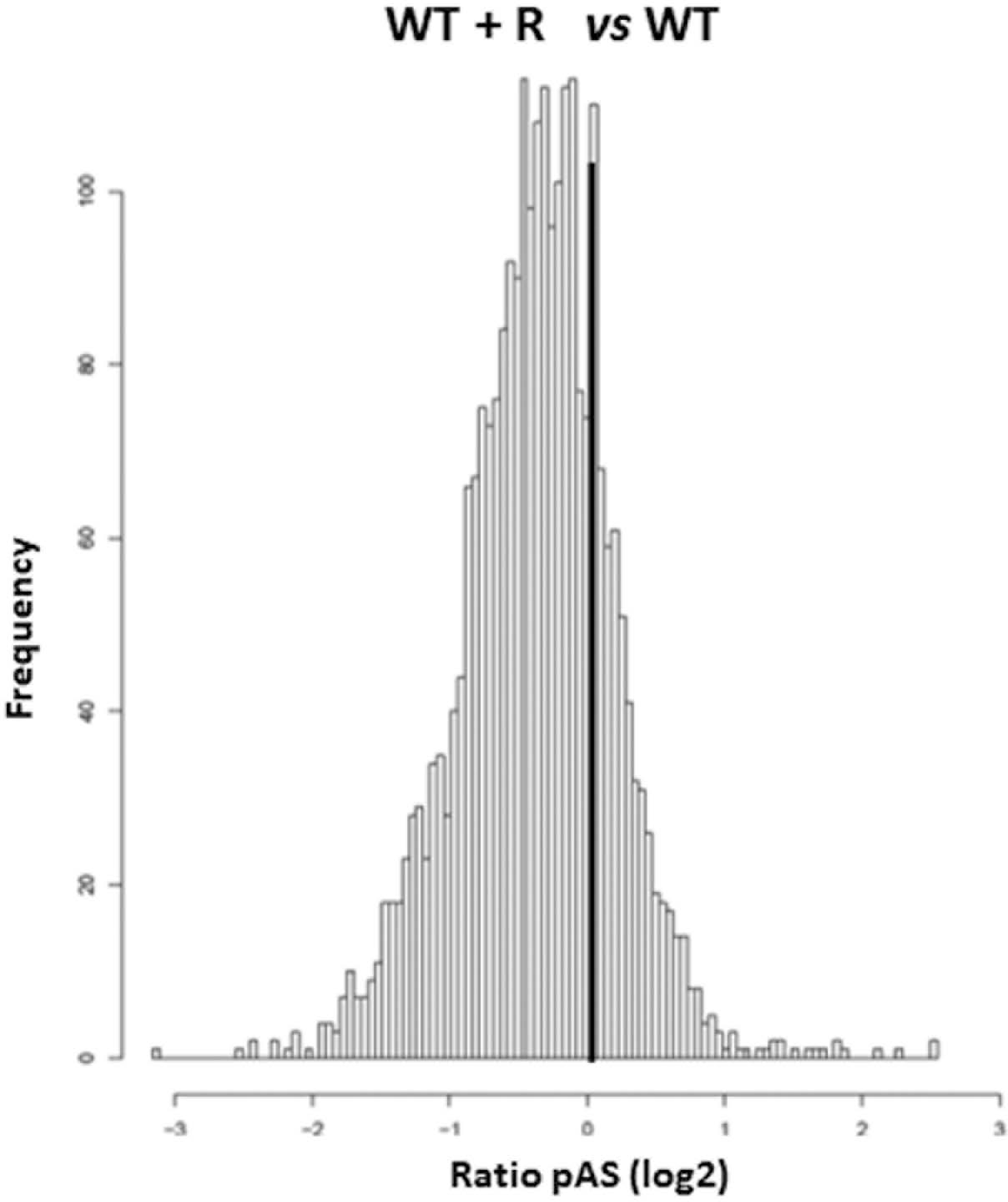
Inactivation of TOR signaling leads to a shift towards upstream polyA sites selection. Gene-specific frequency to preferentially use pAS upstream or downstream of the canonical annotated pA site. The X-axis shows the ratio Log_2_(reads WT+R _down_/ reads WT+R _up_) - Log_2_(reads WT _down_/ reads *bud27Δ*_up_), where a negative value represents preferential upstream pAS use in the presence of rapamycin (WT+R). The Y-axis shows the frequency of the mRNAs corresponding to each value on the X-axis, for wild-type (WT) cells grown with rapamycin (WT + R) to inhibit the TOR cascade *vs.* the cells grown without rapamycin.

It has been reported that inactivating the TOR pathway by rapamycin addition leads to polyA tail shortening and decreased mRNA stability (68). In addition, alteration in alternative polyadenylation and chromatin structure around pAS also affects mRNA stability (22). To investigate whether lack of Bud27 could affect mRNA half-lives, we analysed mRNA decay for some genes. For this purpose, the wild-type and *bud27Δ* mutant cells were treated with 5 μg/ml of thiolutin to block transcription, and, as previously reported, total RNA was extracted at different times after thiolutin addition (69). We analysed mRNA decay for *CPA2*, *HHT1*, *RPB6* and *ACT1* by RT-qPCR and demonstrated diminished mRNA stability in the *bud27Δ* mutant *versus* the wild-type strain (Figure 7). Furthermore, global mRNA decay, which was determined by a dot-blot experiment, also showed that lack of Bud27 globally decreased mRNA stability (Figure 7).

**Figure 7:**
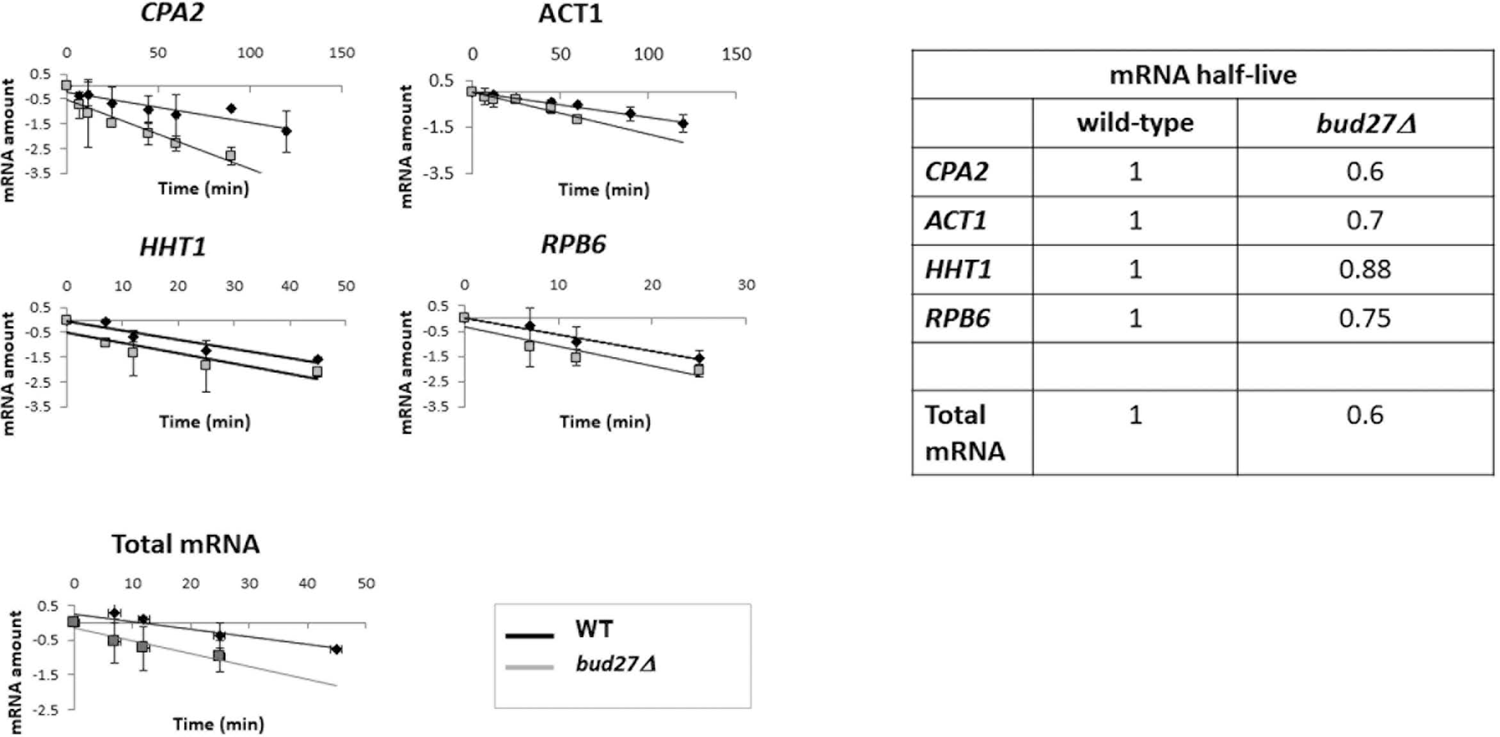
mRNA stability. mRNA levels, measured by RT-qPCR, for genes *CPA2*, *ACT1*, *HHT1*, and *RPB6* at different times after thiolutin addition to block transcription in the wild-type (WT) cells grown at 30°C in both the absence and presence of rapamycin for 12 h to inactivate the TOR pathway. rRNA 18S was used as a normalizer in RT-qPCR. Global mRNA levels measured at different times after thiolutin addition to block transcription in the wild-type (WT) cells with or without rapamycin addition using an oligodT probe by a dot-blot assay. Experiments corresponded to at least three independent biological replicates.

According to these results, the striking similarity between the effects caused by inhibition of TOR signaling on chromatin structure, pAS selection and mRNA stability with those caused by lack of Bud27 lead us to speculate that they might be Bud27-dependent.

## DISCUSSION

RNA pol II transcription elongation depends on prefoldin-like Bud27, which associates with the Rpb5 subunit and the chromatin RSC remodeler complex, and mediates the RSC-RNA pol II interaction. By employing a mutant lacking Bud27 that affects transcription elongation by RNA pol II, we herein demonstrated that RSC associated with chromatin throughout the gene body and influenced nucleosome positioning and chromatin structure around pAS by determining pAS usage, and then, alternative polyadenylation.

The data herein presented and our previous results demonstrate a relation between transcription elongation defects and alteration in RSC association with the RNA pol II and the chromatin in *bud27Δ* mutant (34). All this reinforces the role of RSC in transcription elongation and its localisation in transcribed coding sequences (19,30-32,65,67), in addition to its established role in transcription initiation (28,31,32,63,64,70,71). Furthermore, the recruitment of RSC to genes seems to be linked with transcriptional activity because those genes whose expression was most affected by lack of Bud27 (39) were those showing greater RSC occupancy in a wild-type strain(31). These results establish a positive relation between Bud27 and RSC with the transcriptional activity of RNA pol II, in line with Bud27, as well as RSC, occupying highly expressed genes (31,39). This relation may occur *via* the tripartite association between Bud27, the RNA pol common subunit Rpb5 and RSC (34,45), which likely occurs through the Rsc4-Rpb5 interaction (33). Consequently, we speculated that lack of Bud27 would alter the tripartite Bud27-RSC-Rpb5 interaction, affecting transcription elongation.

Consistently with the reduced elongation rate, we observed an overall increase in RNA pol II occupancy in gene bodies, similarly to that observed upon Sth1 loss (31) or in other RSC mutants (32). As proposed, this could result from the difficulty of RNA pol II to navigate once the RSC-mediated remodeling of the nucleosome is altered, and suggests that RSC might help in RNA pol II elongation by increasing the accessibility of nucleosomal DNA. Nevertheless reduced RNA pol II occupancy in RSC mutants have also been observed, but only for some specific genes (72,73). As a role for Rpb5 in transcription elongation has been reported (13), we cannot rule out that altering the tripartite Bud27-Rpb5-RSC interaction would influence transcription elongation by impacting the RSC association with both RNA pol II and chromatin.

Our data from MNase-Seq and the analysis of histone H3 occupancy showed an alteration in nucleosome positioning and occupancy along gene bodies. These results agree with other previously reported ones obtained upon loss or depletion of RSC subunits, and suggest that RSC is involved in remodeling and sliding nucleosomes and in histone eviction during transcription (28,30–32,63–65,67,70,74–77). Notably, the increase in histone H3 occupancy upon Sth1 loss has also been observed for RNA pol III genes, which supports our data (67). It was noteworthy that the decreased Sth1 occupancy in the 3’ region of gene bodies observed with lack of Bud27 led to a clear alteration to chromatin structure around pAS. This phenomenon has not been previously reported for the RSC mutant or RSC alterations. It is accompanied by greater and differential RNA pol II occupancy both upstream and downstream of pAS, which could account for paused or slower RNA pol II. Furthermore, the differential RNA pol II accumulation downstream of pAS in the *bud27Δ* mutant in relation to wild-type cells could hint at a defect in transcription termination because proper nucleosome positioning at the end of genes has been proposed as being important for transcription termination (78,79). In line with this, a role of RSC in transcription termination or RNA pol II recycling has been put forward because Rsc8-depleted cells accumulate RNA pol II downstream of the TTS (32). Under these conditions however, no significant differences in nucleosome occupancy have been observed (32).

These data suggest that RSC may influence transcription by allowing nucleosomes to be more accessible to elongating RNA pol II and to favour nucleosome remodeling for efficient transcription elongation. In agreement with this, previous evidence suggests that nucleosome remodeling by RSC favours DNA accessibility for RNA pol II transcription (31) and *in vitro* experiments have demonstrated that RSC remodels nucleosome and promotes transcription elongation (29). These results agree with the role of RSC acting on the promoters on the organization of −1 and +1 nucleosomes and, to a lesser extent, to the +2 and +3 nucleosomes (28).

The alteration in chromatin structure around pAS in the *bud27Δ* mutant accompanied by the RNA pol II accumulation, could account for the observed differential pAS usage, which may affect alternative polyadenylation. In agreement with this, the association between alternative polyadenylation and chromatin alteration has been previously established (21–23), and is likely related to histone modifications, as occurs for H3K36me3, H3K4me1 and H3K4me3 (21–24). Interestingly, in relation to histone modifications and/or epigenetic modifications, histone H3 acetylation promotes RSC binding to chromatin (29,30,80). However, this is the first reported data linking RSC with chromatin alteration around pAS and with alternative polyadenylation, and indicates that RSC-dependent nucleosome remodeling which occurs during transcriptional elongation (19,30,31,65,67) also takes place around pAS. Our data show a shift to upstream pAS selection, which agrees with recent data that demonstrate similar behaviour for RNA pol II with a slow elongation rate, while a modest shift to downstream pAS selection is noted for RNA pol II with a fast elongation rate (7,9). Our data also agree with the results obtained in mammals and *Drosophila* containing RNA pol II with slow elongation rate (10,11).

Our data also point out the alteration of alternative polyadenylation upon TOR signalling cascade inhibition by adding rapamycin, similarly to what occurs in response to environmental conditions in yeast (22,81–83), or by environmental or developmental conditions in mammal cells (84-86,87 In line with this, inhibiting TOR with rapamycin in yeast affects alternative polyadenylation, a phenomenon that is associated with epigenetic modifications {Kaczmarek Michaels, 2020 #1983). In this scenario, a drop in the histone H3 levels around pAS occurs (22), while we evidenced a general increase in histone H3 along gene bodies. The fact that Sth1 association with chromatin, Rpb1 occupancy, histone H3 occupancy and pAS usage observed upon TOR inhibition by rapamycin addition correlates with the data obtained when Bud27 is lacking suggest that these phenomena are Bud27-dependent, in line with the participation of Bud27 in the TOR cascade and in the TOR-transcription interconnection (39,40,48). Note that Bud27 human orthologue URI also participates in the TOR cascade (40). Furthermore, the decrease in mRNA stability in the *bud27Δ* mutant strain also agrees with mRNA instability observed in yeast cells upon rapamycin addition, which has been reported to be associated with polyA tail shortening (68). Notably, tail length and mRNA stability have been reported to be related to alternative polyadenylation (87,88).

When collectively taking all this information, we propose that RSC is necessary for maintaining correct chromatin structure during transcription elongation. The modification in chromatin structure provoked by altering the association of RSC with both chromatin and RNA pol II may lead to arrested or slower RNA pol II around pAS, which may enable less used pAS to be exposed. Concomitantly, 3’ processing machinery may easily access these less used pAS by adding the polyA tail. To support this proposal, RSC subunits have been shown to physically interact with polyA machinery elements (https://www.yeastgenome.org/) and similar scenarios have been proposed (7,22). Finally, it would be interesting to investigate whether RSC influences chromatin structure for proper transcription termination.

## ACKNOWLEDGEMENTS

We thank Dr. J.E. Pérez-Ortín for helpful discussion and the “Servicios Centrales de Apoyo a la Investigación (SCAI)” of University of Jaen for technical support.

---

**Figure S1:**
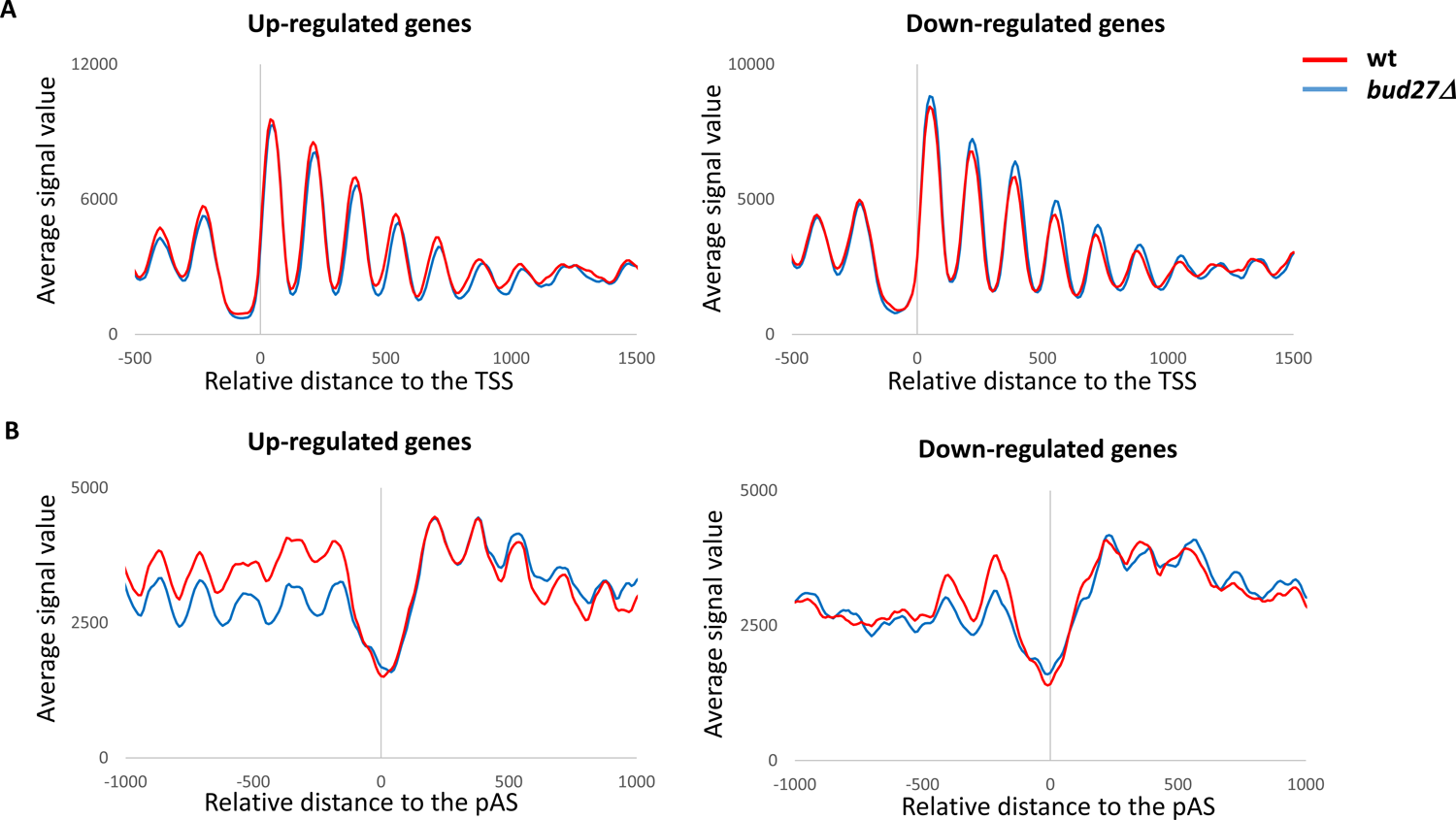
Lack of Bud27 alters nucleosome positioning. Metagene analysis of nucleosome occupancy for the wild-type (red) and *bud27Δ* cells (blue) in the up-regulated and down-regulated genes in the *bud27Δ* mutant *vs.* the wild-type (WT), according to (39). Genes were scaled to the same length and then aligned to their TSS (**A**) or their pAS (**B**). All the genes in the yeast genome, for which a TSS/pAS is available, were considered.

**Figure S2:**
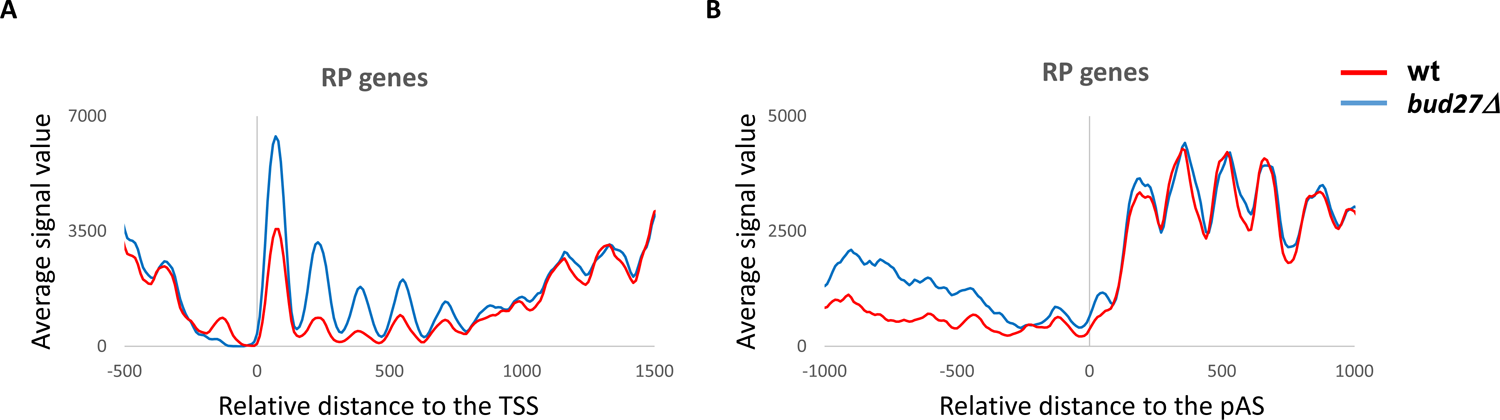
Lack of Bud27 alters nucleosome positioning. Metagene analysis of nucleosome occupancy for the wild-type (red) and *bud27Δ* cells (blue) in RP genes. Genes were scaled to the same length and then aligned to their TSS (**A**) or their pAS (**B**). All the genes in the yeast genome, for which a TSS/pAS is available, were considered.

**Figure S3:**
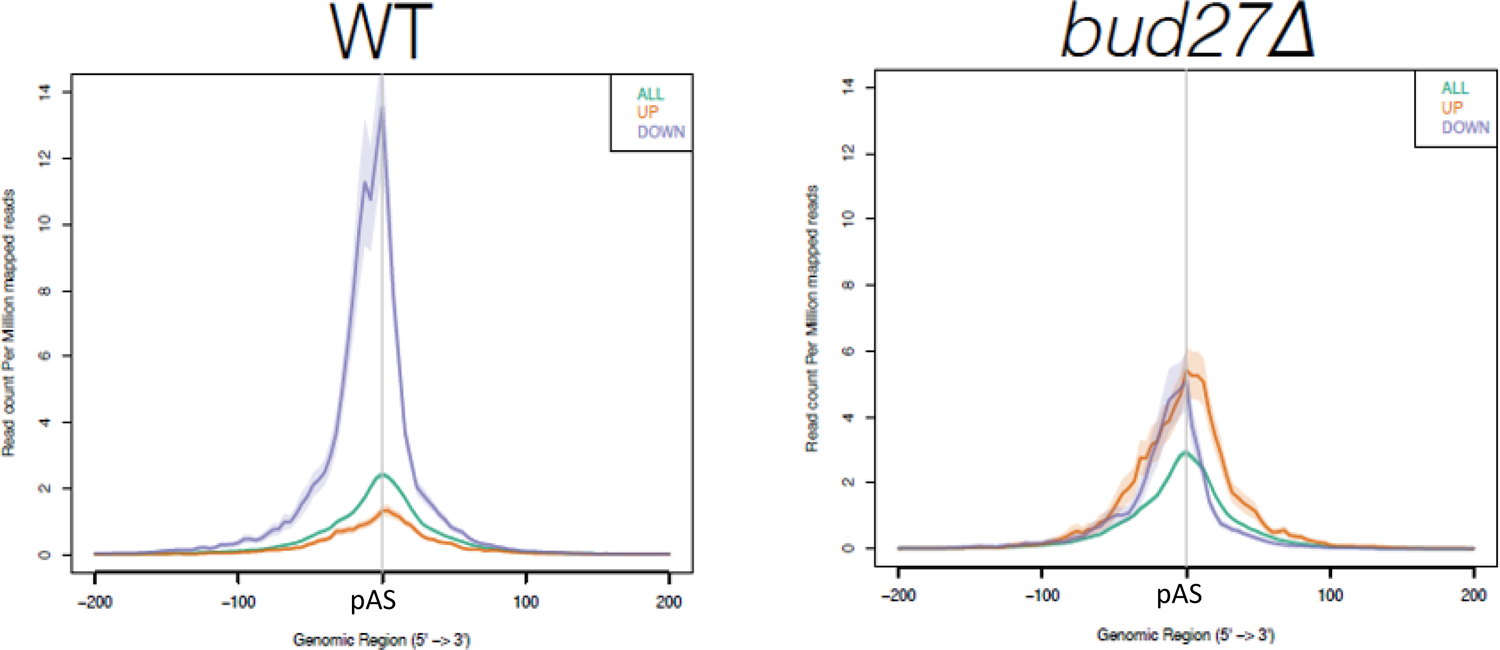
Polyadenylation patterns in wild-type and *bud27Δ* mutant cells. Polyadenylation in the wild-type (WT) and *bud27Δ* cells in the −200 to 200 region around pAS for the up- and down-regulated genes in *bud27Δ* cells, and for the whole set of genes (39).

**Figure S4:**
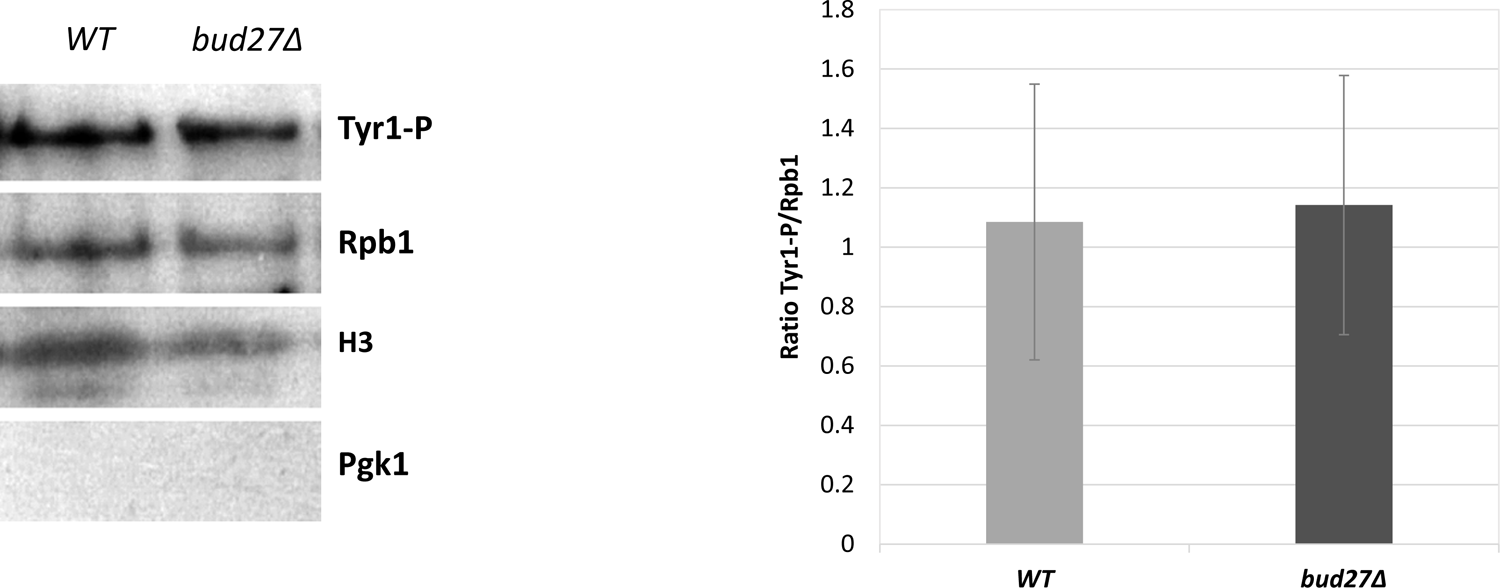
Tyr1 phosphorylation of RNA pol II in the wild-type and *bud27Δ* mutant cells. Western blot with antibodies against Rpb1 and Rpb1-CTD Tyr1-P in chromatin enriched-fractions of the wild-type (WT) and *bud27Δ* cells isolated by the yChEFs procedure (55,89). H3 histone was used as a positive control of chromatin associated proteins, and Pgk1 was the negative control of cytoplasmic contamination. Experiments corresponded to at least three independent biological replicates.

**Supplementary Table S1.**
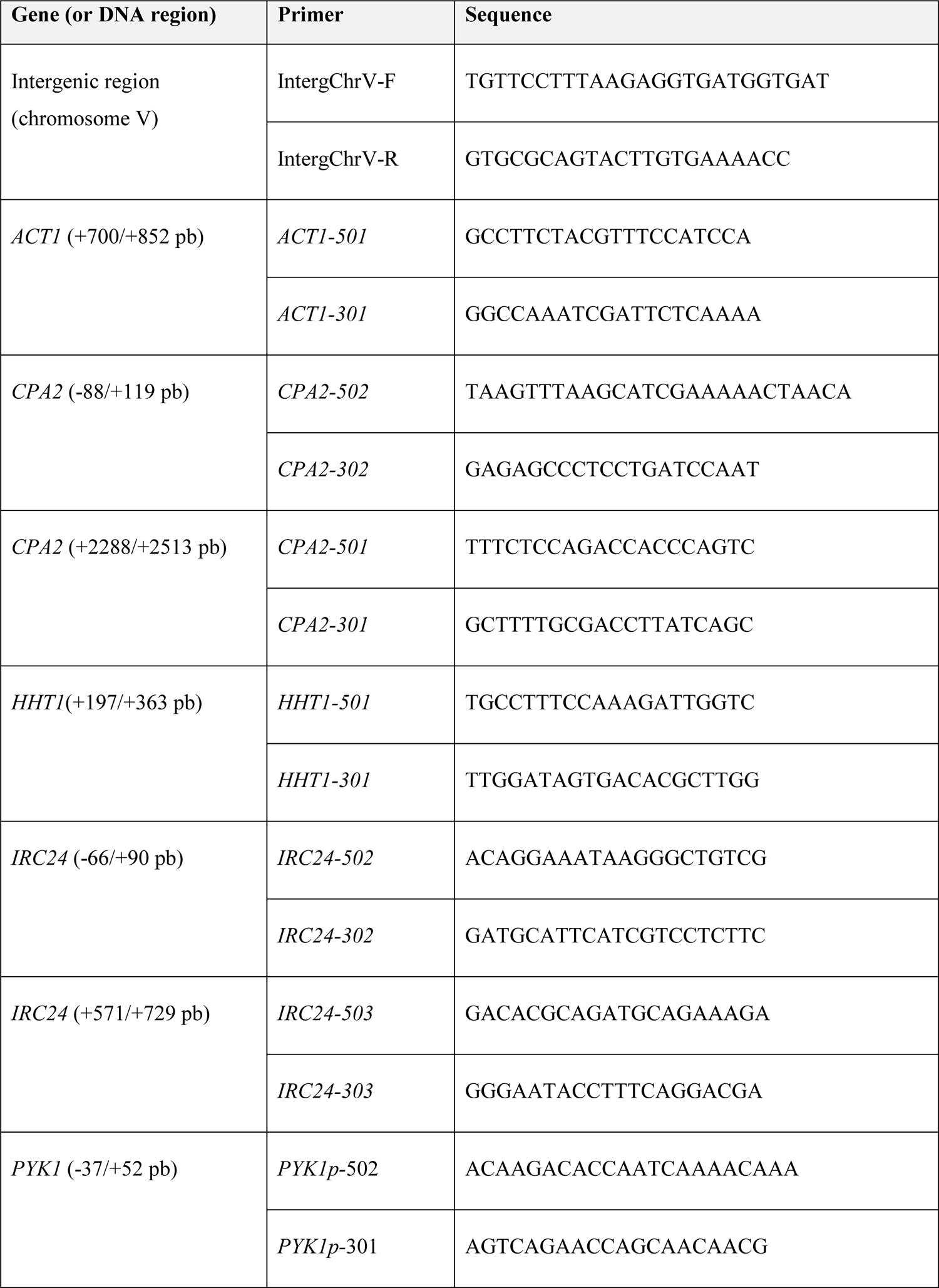
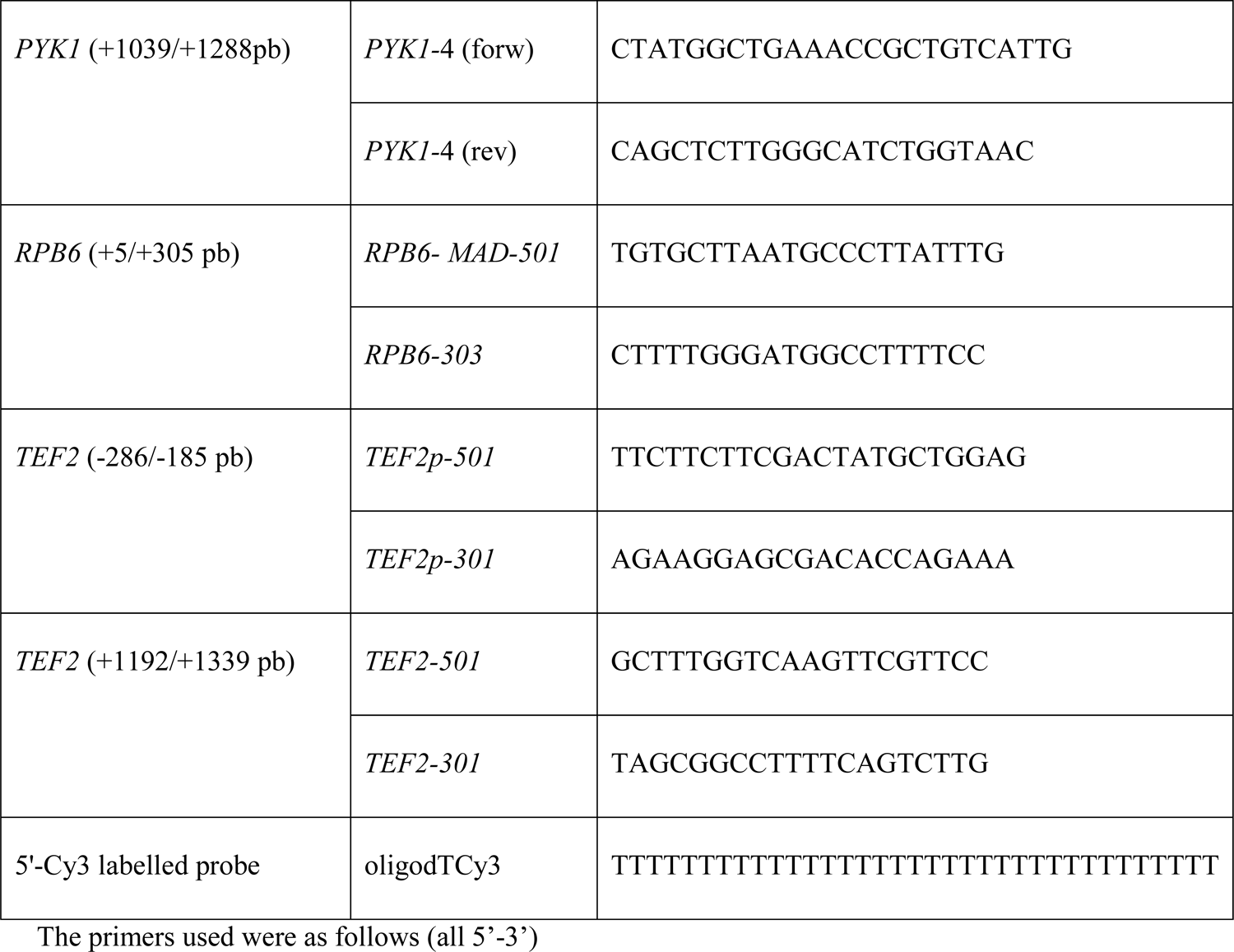
Oligonucleotides used

